# The ectomycorrhizal community of urban linden trees in Gdańsk, Poland

**DOI:** 10.1101/2020.07.30.228668

**Authors:** Jacek Olchowik, Marzena Suchocka, Paweł Jankowski, Tadeusz Malewski, Dorota Hilszczańska

## Abstract

The linden tree (*Tilia* spp.) is a popular tree for landscaping and urban environments in central and northwest European countries, and it is one of the most popular in cities in Poland. Ectomycorrhizal fungi form a symbiosis with many urban tree species and protect the host plant from heavy metals and against salinity. The aim of this study was to characterize the ECM fungal community of urban linden trees along the tree damage gradient. The study was performed on two homogeneous sites located in the centre of the city of Gdańsk, in northern Poland. The vitality assessment of urban linden trees was made according to Roloff’s classification. Tree damage classes were related to soil characteristics using principal component analysis. The five ectomycorrhizal fungal species were shared among all four tree damage classes, and *Cenococcum geophilum* was found to be the most abundant and frequent ectomycorrhizal fungal species in each class. Park soil had significantly lower pH and Na, Cl and Pb content than street soils. Our knowledge of ectomycorrhizal communities in urban areas is still limited, and these findings provide new insights into ectomycorrhizal distribution patterns in urban areas.

## Introduction

The most heavily human-modified ecosystems, cities, are expanding rapidly [1]. Trees provide benefits, ensuring that sustainable urban development and environmental benefits are most valued by city managers as a reason for introducing trees into cities [2]. Nevertheless, paradoxically, they grow in often extremely distorted habitat conditions in comparison to natural conditions. Street trees are exposed to a relatively high stress level. Studies reveal that their average lifespan is shorter than that of park trees [3], with mean ranging from 19 to 28 years [4] or less. The linden tree (*Tilia* spp.) is a popular tree for landscaping and urban environments in central and northwest European countries [5] and it is one of the most popular in cities in Poland. Dmuchowski and Badurek [6] reported that in Warsaw during 1973-2000 over 50% of trees growing alongside the four main thoroughfares in the city centre were removed. Moreover, the continuation of these studies has shown that over a period of 35 years, out of the 5 species with the highest loss, 3 were *Tilia* species: *Tilia platyphyllos, Tilia ‘Euchlora’* and *Tilia cordata* [7]. The stresses that affect urban trees may be biotic or abiotic, mechanical damage, high temperature, soil compaction, limited soil volume for root development and drought [8,9,3]. Specifically, soil and roots may be affected by construction activities such as utility trenching, soil compaction and subsequent root deoxygenation, shortage of available water, and incorporation of anthropic materials [5,10,11]. Under stress, plant growth and photosynthesis are reduced and carbon allocation is altered, resulting in a low tree vitality [12,13,14,15,16,17].

Mycorrhiza is a mutualistic association because fungi form relationships in and on the roots of a host plant. Mycorrhizae protect the host plant from heavy metals and against drought [18]. Ectomycorrhizal fungi (ECM) are ecologically significant because they provide the plant with several benefits including enhanced nutrients [18] and increased water use efficiency, and enhanced root exploration. Mycorrhizal colonization has been shown to promote short root survival particularly when *Tilia* trees are exposed to drought conditions. Ectomycorrhizal fungi promote water uptake in general [e.g., 19] and have been specifically shown to play an important role in the nutrient uptake of *Tilia* spp. [20]. Mycorrhizae protect the host plant from heavy metals and promote short root survival particularly when *Tilia* trees are exposed to drought conditions [18,12]. It has been reported that this symbiosis plays a major role by increasing the efficiency of sodium-excluding mechanisms in infected roots and through higher root accumulation of phosphorus (P) [21]. The ecological distribution of fungi is markedly different from natural and urban environments, where mycorrhizal fungi have evolved and adapted. For example, Timonen and Kauppinen [22] reported that *Tilia cordata* trees growing in a nursery had different sets of ectomycorrhizal symbionts than trees grown along streets with traffic, but still the relationship between specific environmental conditions and the mycorrhizal status of trees is not well known [23].

The aim of this study was to characterize the ECM fungal community of urban linden trees along the tree damage gradient. We hypothesized that the ECM fungal community changes along the damage gradient, as the diversity of ECM fungi increases in accordance with tree health condition.

## Materials and methods

### Study sites

We performed the study on two homogeneous sites located in the centre of the city of Gdańsk, in northern Poland. The study was conducted on trees belonging to one genus, *Tilia*: Dutch Linden (*Tilia x europaea*), Fine Linden (*Tilia cordata*), and Broad-leaved Linden (*Tilia platyphyllos*). The first study area was located in the middle of Great Linden Avenue (54°22′05,5″N 18°37′51,2″E), which is a four-row avenue created in 1768-1770 and located within the administrative borders of the City of Gdańsk. The avenue is located within one of the most important and busiest transport routes in Gdańsk. Great Linden Avenue is subject to legal protection under the Act of 16 April 2004 on Nature Conservation and the Act of 23 July 2003 on Monuments Protection and Care as an object entered in the register of monuments, no. 285 of 23.02.1967. Selecting a linden alley as the research area enabled us to exclude the variable of other trees, particularly deciduous species, affecting the community dynamics of *Tilia*-associated ECM fungi. The second site was located in a park (54°22′07,68″N 18°37′57,36″E) at a distance of approximately 150 m away and separated from the road by a dense strip of bushes and hedges.

### Tree health assessment

The vitality assessment of each tree was made according to Roloff’s classification [24] and the health condition of the trees was estimated according to leaf and branch growth pattern. Condition evaluation was performed for each tree and was based on distal crown vigour. Trees were segregated into 4 groups: R0 ‘exploration’ (tree in the phase of intensive offshoot growth), R1 ‘degeneration’ (tree with slightly delayed offshoot growth), R2 ‘stagnation’ (tree with visibly delayed offshoot growth), R3 ‘resignation’ (tree is dying, without the possibility of regeneration or returning to the second class). In the first study area, thirty street trees at least 200 m apart were classified according to the declining classes and assigned to classes R1, R2 and R3. At the park site ten trees belonging to class RO were selected. All the trees situated along the street and in the park site were of the same age (60 years).

### Sampling and identification of mycorrhizae

In May 2019 soil cores were collected from both street and park trees. For each of 40 trees, a total of 80 soil subsamples were collected for mycorrhizal assessment: each sample consisted of 2 microsite localities: north and south (40 trees × 2 microsite (north and south) = 80 subsamples). The street root samples were taken from the 1.5 m wide grass strip between roadways. To access the root system, each sample was extricated with a cylinder (approximately 25 cm diameter, 25 cm depth) of adjacent substrate and packed in labelled plastic bags. Samples were stored at – 20 °C until further processing. Extracted root fragments were examined under a dissecting microscope at 10-60x magnification.

All tips were classified as ‘vital ECM’ (VM, with ECM mantle) ‘non-vital’ (NV, scurfy surface, without remnants of ECM mantle) or ‘vital non-ECM’ (NM, well-developed, and mantle lacking) [25]. Mycorrhizae were classified into morphotypes based on morphological characters (colour, shape, texture, and thickness of the mantle, presence and organization of the emanating hyphae, rhizomorphs, and other elements) according to Agerer [26], and the experience of the researchers involved in this study [27]. The degree of mycorrhization of linden roots, abundance, relative abundance and frequency of individual ectomycorrhizal fungal taxa were determined according to Olchowik et al. [27]. Each morphotype was treated separately during molecular identification and was pooled for calculation of abundance only after molecular analysis indicated that morphotypes belonged to the same taxa. The internal transcribed spacer (ITS) region of the rDNA was amplified using the primers ITS1F and ITS4 [28,29] and the product of the polymerase chain reaction (PCR) was sequenced. The full methods used for molecular identification of mycorrhizae are reported by Olchowik et al. [30]. The best representatives of each unique ITS sequences were deposited in NCBI GenBank with the accession numbers.

### Physicochemical analysis of the soil

The samples for soil chemical analysis were taken at the beginning of May 2019. The samples were collected from 30 trees in the alley and from the park area, which included 10 trees growing in the neighbouring area, within the boundaries of the city park. Samples of soil were air-dried, passed through a mesh screen, and stored for further analysis. The soil analyses were performed in the laboratory of the Polish Centre for Accreditation (No. AB312). The accuracy of the analysis was checked against standard reference materials: international standard soils [31-34]. The phosphorus (P) was determined for all samples with 1% citric acid extraction, according to Schlichting et al. [35]. The soil pH and was determined by mixing 20 ml of soil substrate with 40 ml of deionized water measured with a calibrated pH meter equipped with a glass electrode.

### Data analysis

For the purpose of data analysis, the two mycorrhizal data subsamples were summed for each tree in order to match the number of soil samples. Hence, a total of 40 samples were analysed in the study. All soil characteristics which were measured below the limit of detection were substituted with the half value of the corresponding limit. In order to investigate the relation between the tree damage classes and the abundance of VM, NM, and NV root tips, the date were cross-tabulated into a contingency table and the chi-square test of independence was performed. The cells in the contingency table which were responsible for the significant departure from independence of the examined variables were identified as those for which the absolute maximum of Pearson’s residual exceeded the value of 2.

The species diversity for each class of trees was estimated with the Chao1 and Shannon diversity indices. The differences in the characteristics of the soil samples between the tree classes were examined with the one-way analysis of variance (ANOVA) or the non-parametric Kruskal-Wallis test. The Kruskal-Wallis analysis was applied in the case of the soil parameters which did not fulfil the assumptions of the ANOVA: the homogeneity of variance (Levene’s test) and/or normality (Shapiro-Wilk test). In the case of significant differences, Tukey’s honestly significant difference (HSD) test (for ANOVA) and Dunn’s test (for Kruskal-Wallis) were used to identify the homogeneous groups of tree classes. Spearman correlation and principal component analysis (PCA) were used to relate the soil characteristics with the abundance of VM, NM, and NV root tips. The Kaiser-Meyer-Olkin (KMO) measure of sampling adequacy was applied to select the variables applicable for the PCA with the KMO threshold value equal to 0.6. Bartlett’s sphericity test was then used to confirm that the set of selected variables is suitable for structure detection.

## Results

The smallest degree of mycorrhization was observed in class R3 (18%). From mycorrhizal root tips after regrouping and combining on the basis of the results of the molecular analysis, finally 11 fungal taxa were detected and assigned to a species level (Table 1, Fig 1). The five ECM fungal species (*Tylospora asterophora* (Bonord.) Donk, *Inocybe grammopodia* Malençon, *Inocybe pelargonium* Kühner, *Cenococcum geophilum* Fr., *Tuber rufum* Picco) were shared among all four trees damage classes (Table 1). *Cenococcum geophilum* was found to be the most abundant and frequent ECM fungal species among all classes (Fig 2a). Moreover, *C. geophilum* was present in more than 60% of all ECM tips (Fig 2b). For each damage class, the species composition of ECM fungi, fungal species richness and diversity indexes were analysed. The number of observed root tips decreased, from the park trees, through the successive tree groups of increasing level of damage. Taxa richness decreased similarly. The numbers of observed root tips in trees from the R0, R1, R2 and R3 groups were 5956, 4472, 3022 and 1163, and the numbers of species were 10, 8, 7 and 5, respectively. For the individual trees, the numbers of observed root tips and the numbers of species were highly correlated: Spearman correlation equal to 0.77 at p-value<0.0001. Due to lack of singletons and doubletons observed in the analysed samples, the Chao1 index computed for each tree class equalled the taxa richness.

**Table 1.**
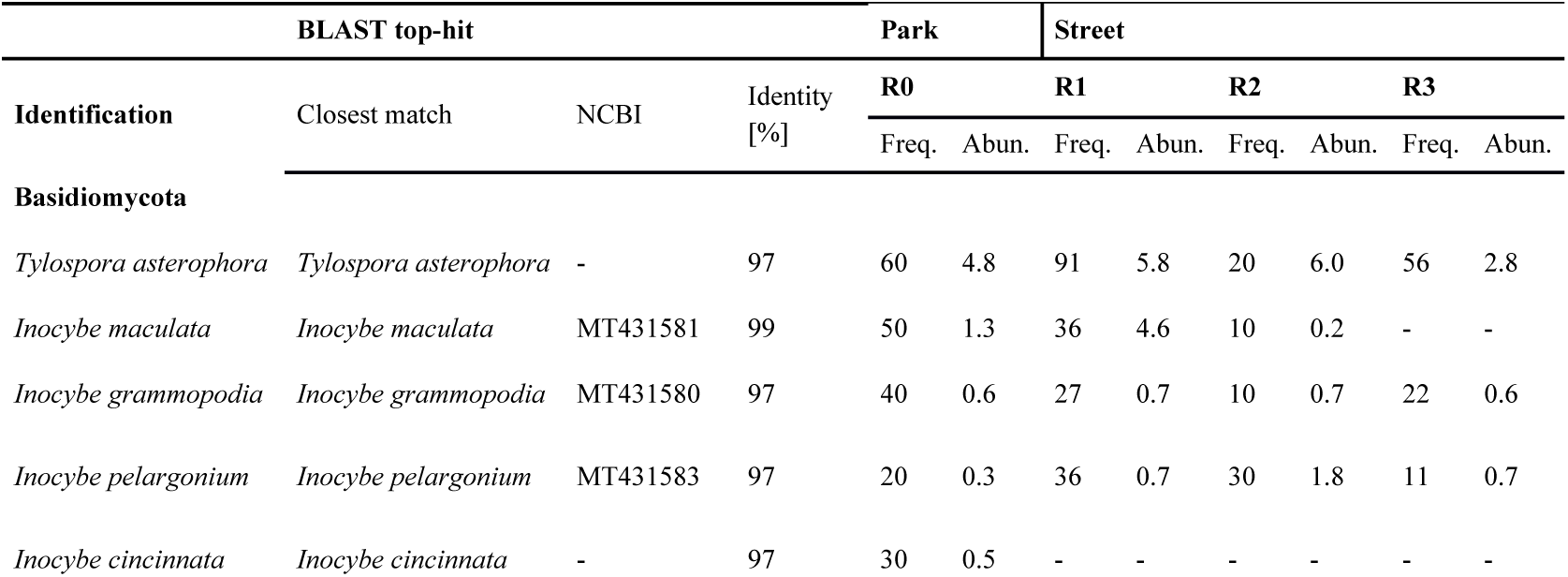
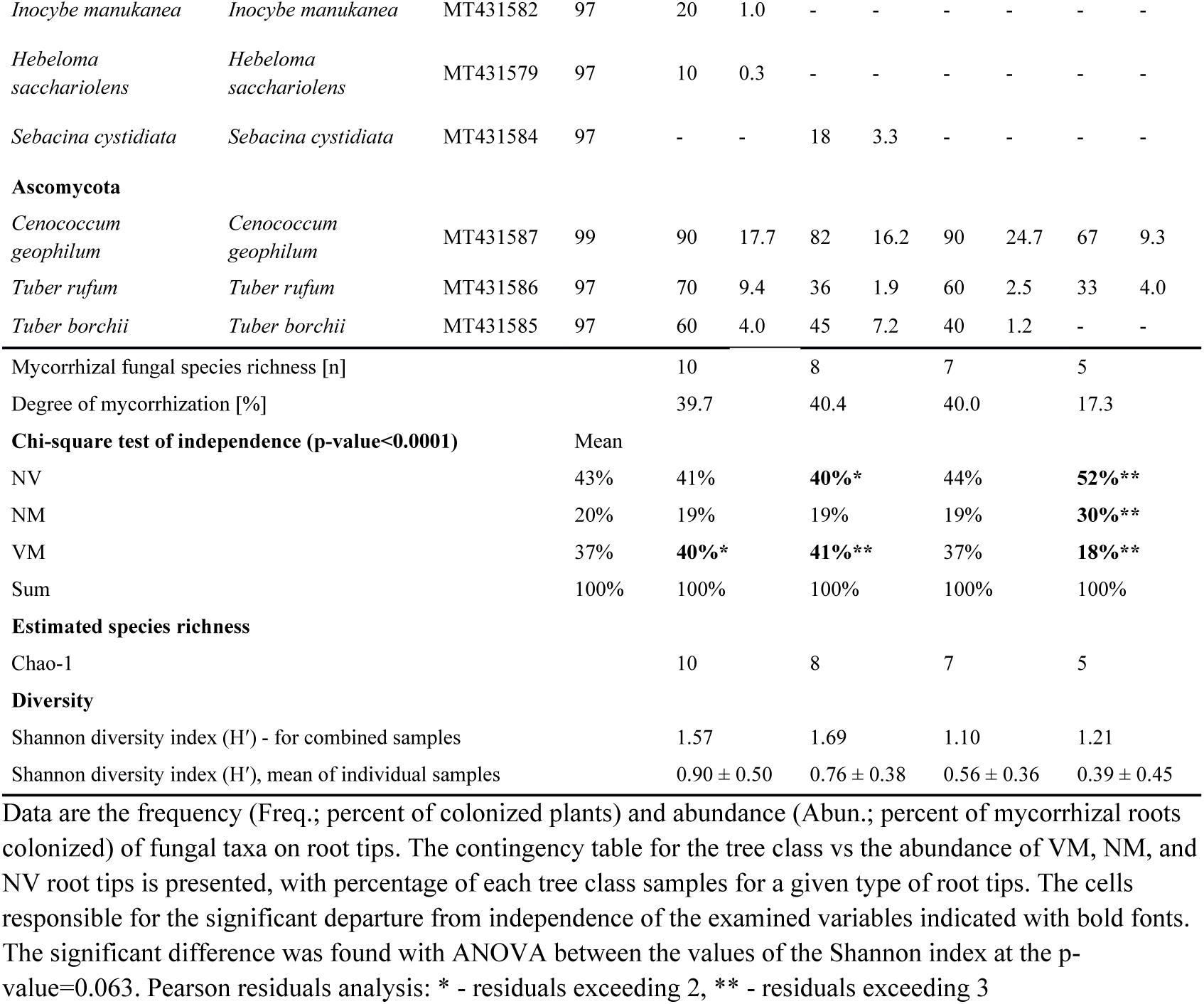
Estimated species richness, diversity and occurrence of fungal taxa associated with the roots of linden trees.

**Fig 1.**
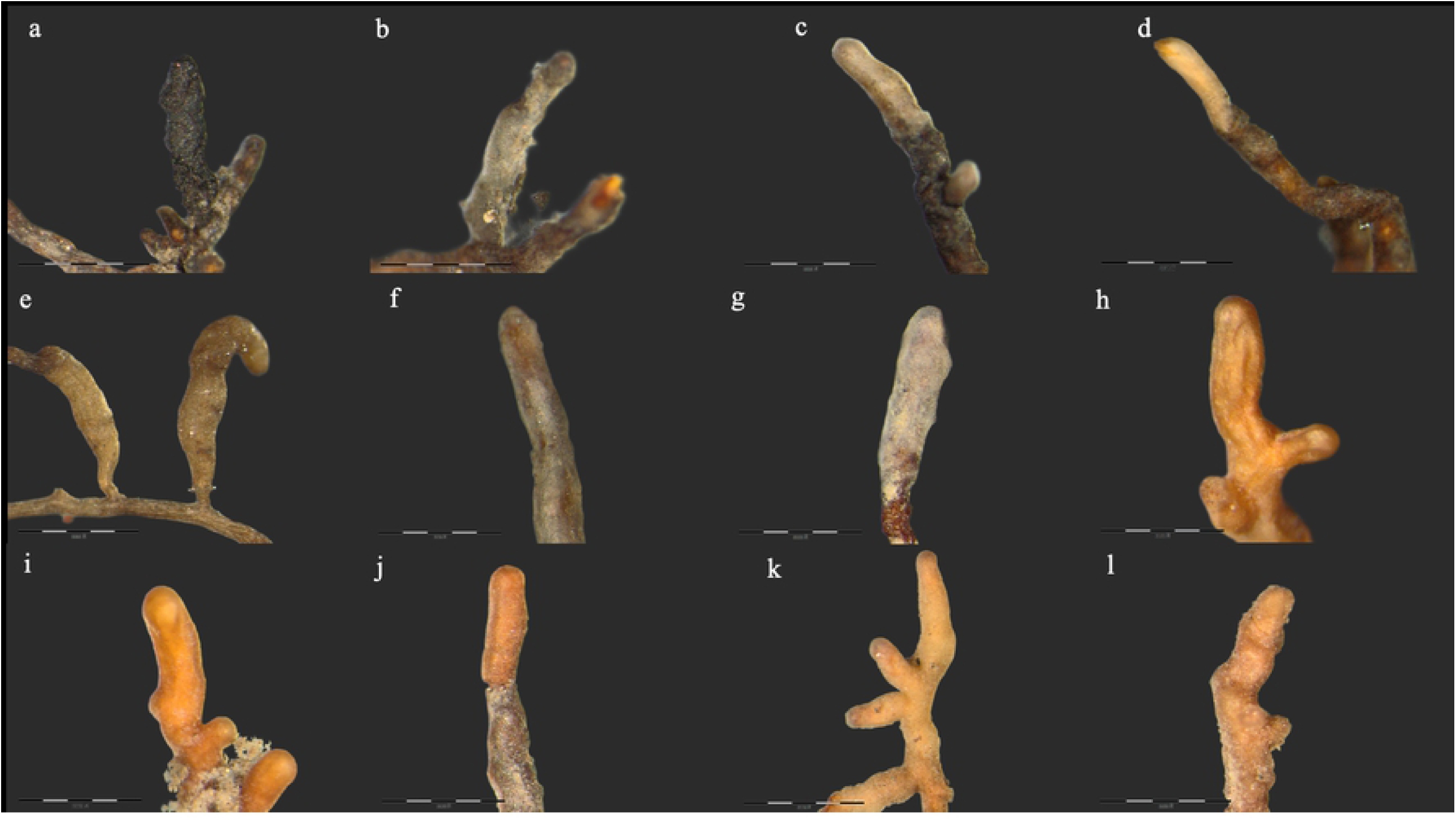
Ectomycorrhizas observed on linden trees in the Gdańsk. (a) *Cenococcum geophilum* Fr., (b) *Hebeloma sacchariolens* Quél., (c) *Inocybe cincinnata* (Fr.) Quél., (d) *Inocybe grammopodia* Malençon, (e) *Inocybe maculata* Boud., (f) *Inocybe* manukanea (E. Horak) Garrido, (g) *Inocybe pelargonium* Kühner, (h) *Sebacina cystidiata* Oberw., Garnica & K. Riess., (i) *Tuber borchii* Vittad., (j) *Tuber rufum* Pollini, (k, l) *Tylospora asterophora* (Bonord.) Donk. Bars in each photograph indicate 0.4 mm length.

**Fig 2a.**
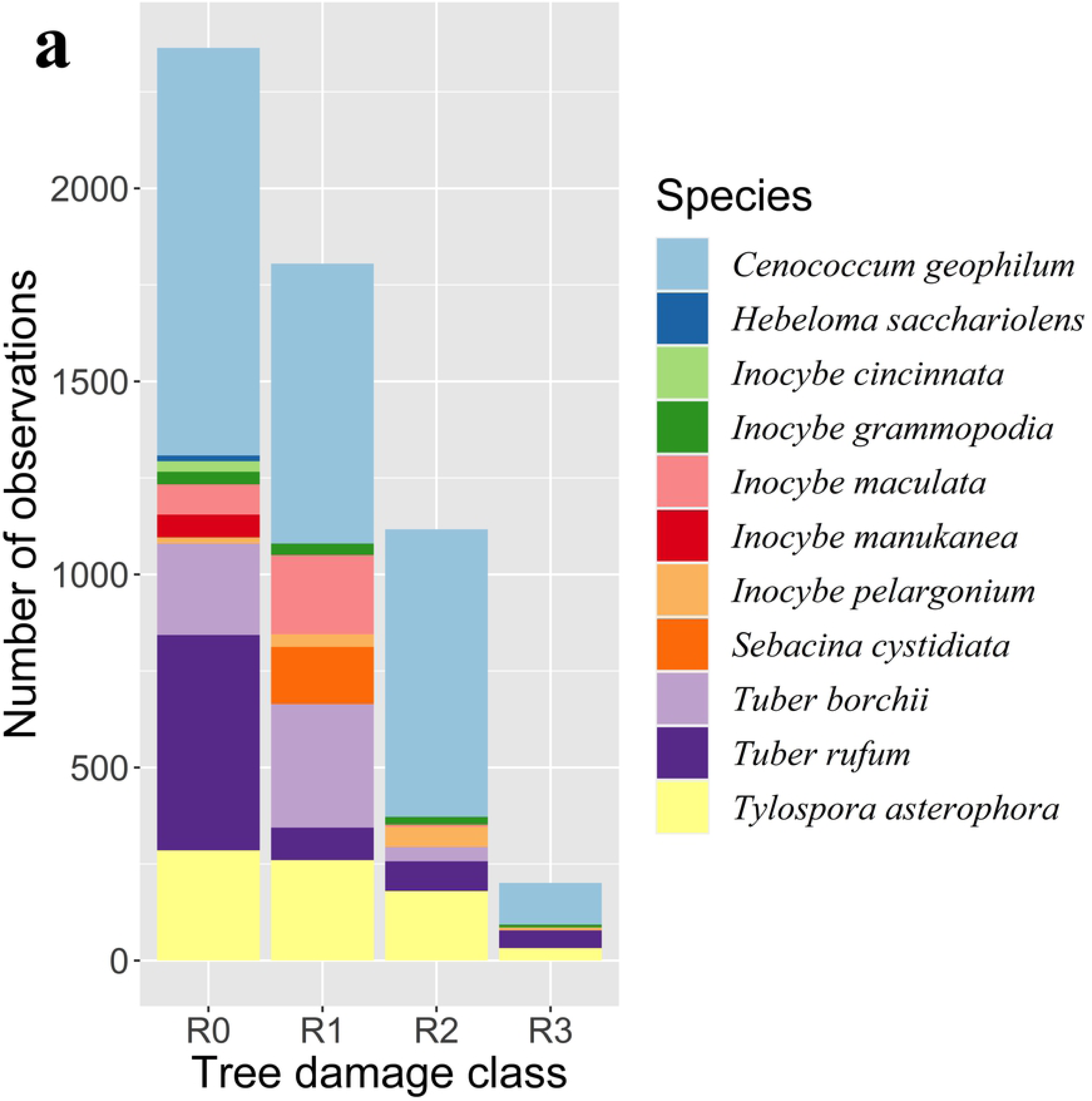
The abundance of the observed ECM fungi species in different damage classes. Each colour represents the number of the root tips with the observed fungi species.

**Fig 2b.**
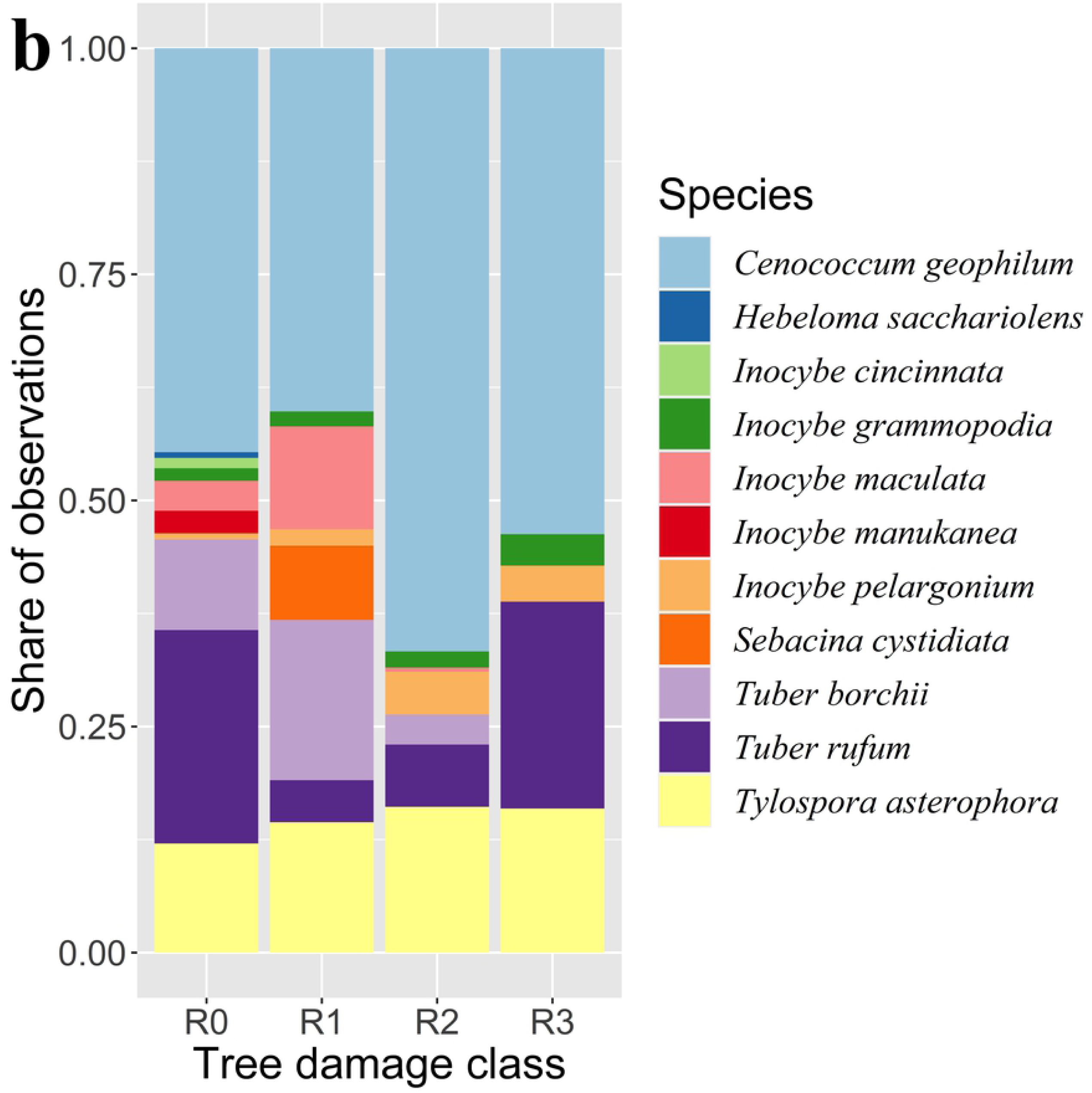
The percentage of the observed ECM fungi species in different damage classes. Each colour represents the number of the root tips with the observed fungi species.

Nearly half of the tested tip samples, 43%, were non-vital, while 20% and 37% of the samples belonged to the NM and VM types (Table 1). The chi-square test showed that, in comparison to this average distribution of the tip classes, the park trees showed a slight excess of the VM type tips, street trees from the R1 group showed an excess of the NV and VM type tips, and samples from the R3 trees had strong overrepresentation of the NV and NM tips and underrepresentation of the VM class tips.

Mean values of the soil parameters between classes are reported in Table 2. In the case of 6 soil characteristics, out of 16 examined, significant differences were found. These parameters were Cl, Na, Pb, Ca and Fe contents and the soil pH. Park soil had significantly lower pH and Na, Cl and Pb content than street soils. Considerable differences were observed between the content of Ca and Fe. In the first case, there was a significant difference only between the trees from the R0 and R3 damage classes, with the park trees having the lowest Ca content. In the case of Fe, there was a significant difference between the trees from the R0, R1 and R3 damage classes, with the park trees having the highest Fe amount. Also, in this case the average Fe content in the samples from the R2 tree class was much lower than in the case of the park trees, but no significant differences were reported due to large variability of the R2 samples. In some cases (Cr for example), though the differences between means seem large, no significant differences were found due to high variability of the data, especially among the R2 tree class samples.

**Table 2.**
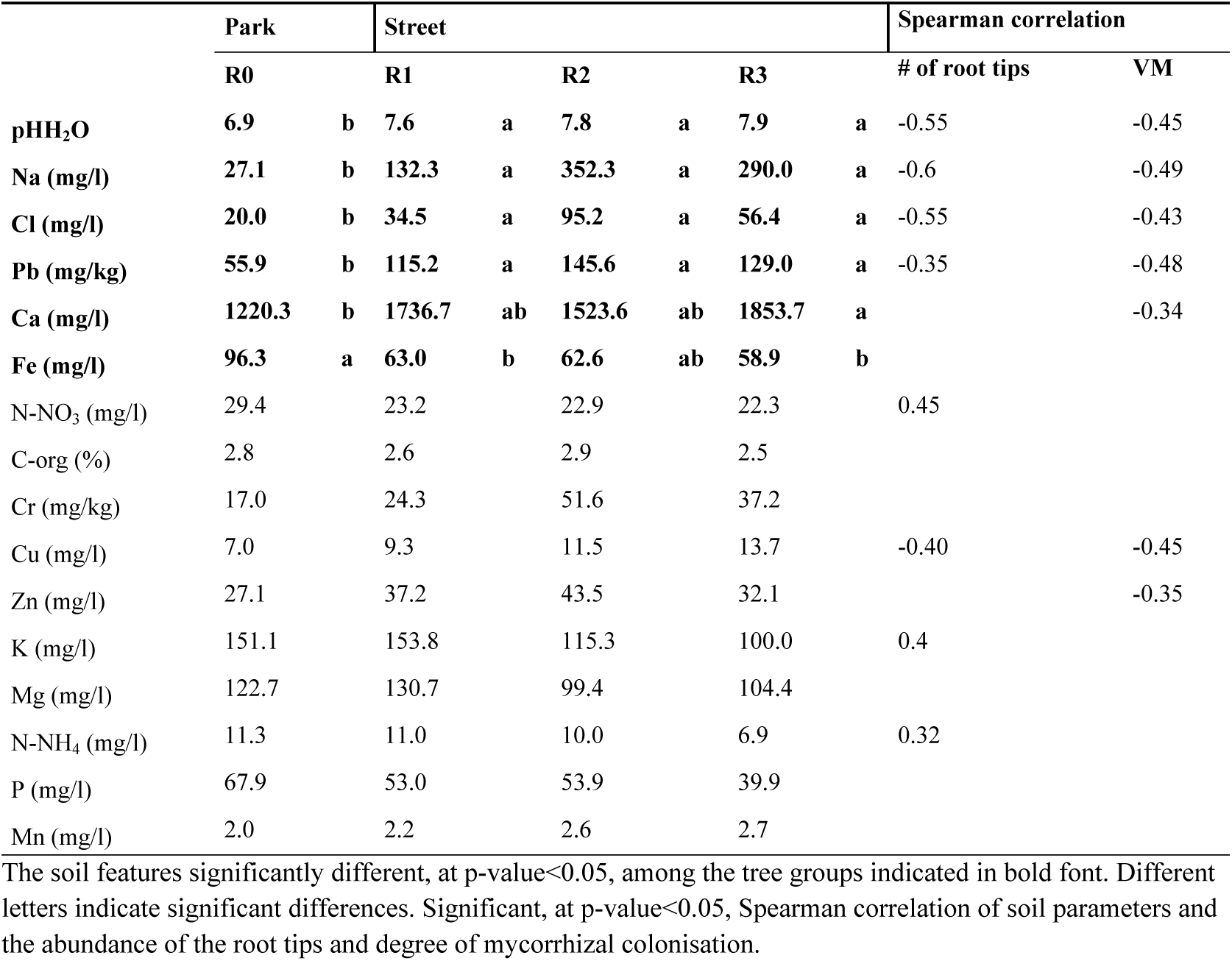
Mean values of selected physical and chemical properties of soil in samples associated with the trees of different damage classes.

Only some of the examined soil parameters were related to the abundance of the root tips and degree of mycorrhizal colonisation (Table 2). As can be seen, increase of three soil parameters, N-NO_3_, N-NH_4_ and K, leads to an increase of the number of root tips. All the remaining soil features negatively influence the abundance of the root tips and the relative abundance of the mycorrhizal root tips (VM).

The overall high variability of the soil characteristics in the samples related to the individual trees can be seen in the PCA plot in Fig 3. Correlation of the examined soil parameters allowed two main groups of them to be distinguished. The first group contains C-org, Cl, Cr, Cu, Na, Pb and Zn, and the second contains K, Mg and N-NH_4_. All members of the second group were negatively related to some representatives of the first one, namely Cu, Cr, Na and Pb. The parameter which links the two groups was pHH_2_O, positively correlated with Cu, Cr, Na and Pb and negatively with N-NH_4_. In the case of the park trees, the samples are distributed parallel to the second group of soil parameters – Mg, K and N-NH_4_ – and in the case of the street trees, the high variability. The remaining soil features, Ca, Fe, P and N-NO_3_, had a weaker correlation with other parameters and Mn showed no relation to any parameter.

**Fig 3.**
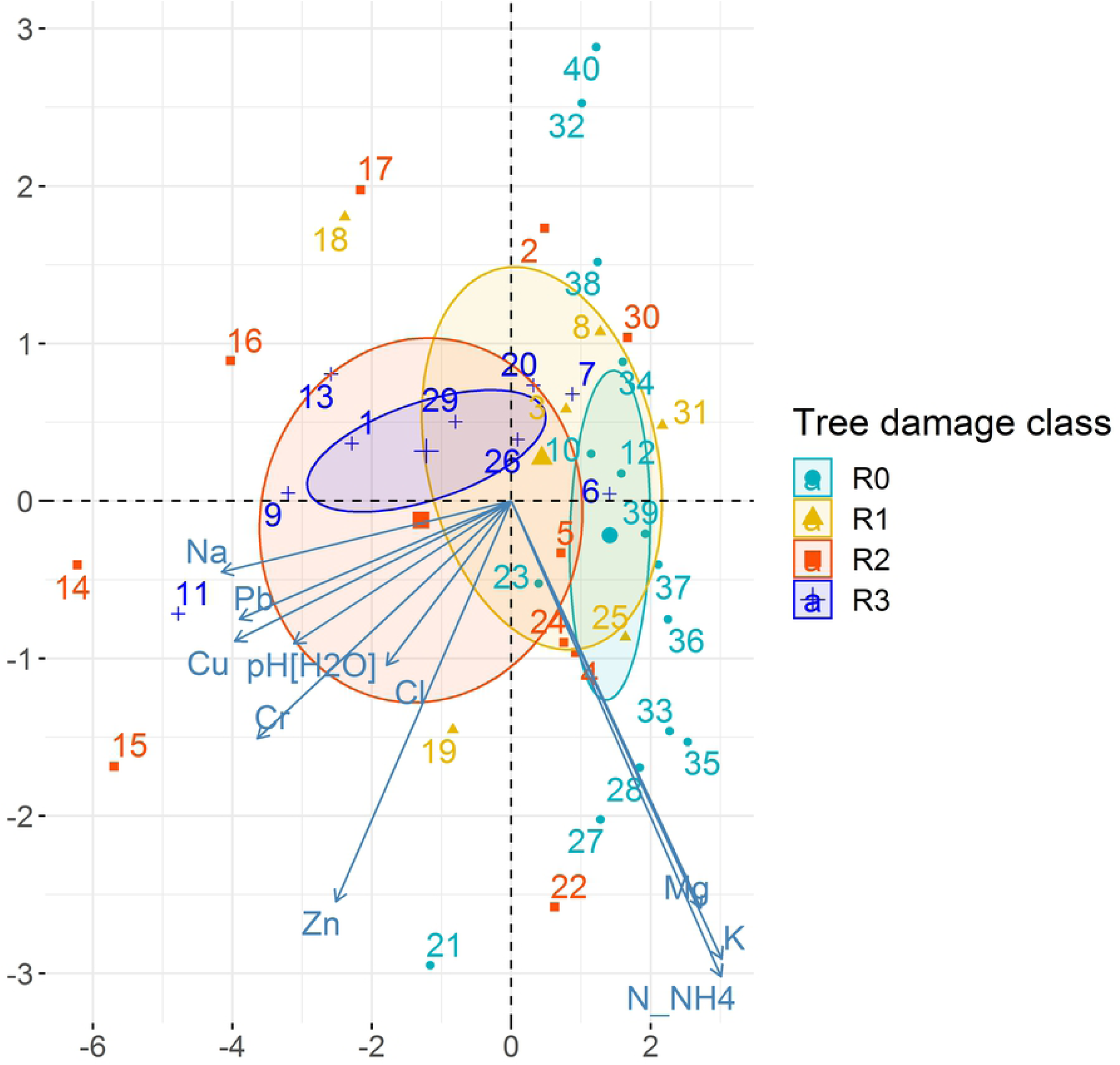
PCA plot representing the relationship between soil parameters studied and their links with classes of trees. The ellipses are the 95% probability confidence ellipses around the mean point of each tree class.

## Discussion

The study presented here investigated the relationships among the health status of urban linden trees using Roloff’s classification with ECM communities. So far, few studies have dealt with the ECM community in urban linden trees [22,36,37,38]. Considering the mycorrhization degree, only class R3 had significantly fewer vital ectomycorrhizal tips than other classes. These data confirmed the data obtained by studying the English oak trees [39], where fine roots of most declining trees had a lower proportion of vital and ectomycorrhizal tips.

The *Tilia* species analysed in our study belong to Great Linden Avenue, which is subject to legal protection. Our study showed that the ECM community structure is highly dependent on the level of decline of the linden trees. The observed ECM fungal species diversity differed significantly along the tree vitality. A similar study conducted in Italy, comparing the health situation of linden trees, classified as ‘moderately declining’ and ‘strongly declining’, showed that the number of ECM fungal species was lower in this second group in comparison to the first [36]. Since lack of nutrients, attack from pathogens, drought and use of de-icing salts are among the main causes of damage of urban trees [40], ECM fungi of urban trees may enhance their growth and survival in the urban environment. In our study the street trees were already 60 years old and the mycorrhizal fungal population associated with their roots was likely to be well adjusted to the street habitat. *Tilia* roots in the park site harboured a diversity of ectomycorrhizal fungi. The number of 10 mycorrhizal morphotypes found in the park in this study was similar to the 12–13 morphotypes observed by Nielsen and Rasmussen [41] in native and planted forests in Denmark. The higher diversity of ectomycorrhizal fungi in the park may be the result of low soil pH in park soils and also partly due to the higher diversity of other ectomycorrhizal plants surrounding the *Tilia* trees in this habitat than in the ‘street’ habitat.

As hypothesized, a gradual increase in taxa richness was observed from the highest damage of trees (R3: 5 taxa) to the best health condition of trees (R0: 10 taxa). The differences among the ECM fungal communities harboured by linden trees on the studied sites may be affected by salinity and concentration of heavy metals. The salt applied to roads in winter is a serious cause of damage of urban trees [42], including water deficit, soil compaction, ion toxicity and ion imbalance [43,44]. Moreover, Na and Cl may inhibit enzymatic activity of fungi [45]. In our study the elevated amount of Na and Cl was the soil feature unique to the street when compared with the park habitat. The soil microbial communities are affected more by salinity than by extremes of any other abiotic factor [46], so this factor could have affected the lower species composition of the ECM fungi associated with the linden street trees.

The PCA analysis showed a gradual shift in similarity between the adjacent damage classes (Fig 3). In part, these differences were due to a significantly higher concentration of heavy metals (Pb, Cr and Cu) in street soil. Moreover, in our study the concentration of Pb in street soil was significantly higher in comparison to park soil. In general, increased concentrations of heavy metals in the soil are known to negatively affect biodiversity [e.g. 47,48]. Heavy metals damage proteins, lipids and DNA [49]. Turpeinen, Kairesalo and Haggblom [50], who investigated the impact of heavy metal contamination on microbial communities, found a negative effect of metal pollution on fungal diversity. This is consistent with the findings in our study, where the street site was shown to host a lower ECM fungal richness than the park site. On the other hand, van Geel et al. [37] reported that the variability in ECM communities of *T. tomentosa* urban trees was little attributed by heavy metal pollution. It is important to note that Van Geel et al. [37] used high-throughput sequencing (HTS) as the basis of taxa identification and the results featured only mycorrhizae identified at the family level. Another point when comparing those results is the different sampling area included in the studies. The study of van Geel et al. [37] was performed on a relatively large scale, due to its location in in three European cities. In our study we concentrated on one city and one street, which limited the potential for replication. More research is needed on a larger sample to reliably identify the reasons for the differences observed between our results and previous research.

Surprisingly, we also found several ECM that were common to all damage classes. There were genera belonging to early-stage fungi, including *I. grammopodia* and *I. pelargonium*. These fungi are often found in habitats with limited nutrient availability [51], for instance in urban ecosystems. Although *T. rufum* needs a more stable habitat [52], this fungus was abundantly present in all damage classes. It may have resulted from the alkaline conditions in street soil, because Tuberaceae generally prefer more alkaline conditions [53]. This result may also suggest that some genotypes are either adapted to street conditions or they are not outcompeted.

The ECM fungal species that we found to be predominant – *C. geophilum* – was present among all damage classes. The dominance of *C. geophilum* was not a surprising result, because this fungus is known as the most efficient drought-tolerant type [12,54]. Considering the ECM community composition related to plant health status, Timonen and Kauppinen [22] demonstrated that *Cenococcum* spp. were more dominant in the roots of unhealthy street trees. This fungus forms ECM with many tree species because of its pioneering capabilities and persistence of sclerotia in the soil [55]. Due to its active growth at low soil temperature and drought tolerance [12] it is well known for its extremely wide habitat range and for being competitive under adverse climate conditions. In the case of our study, abundant root colonization by *C. geophilum* may also be the result of competition for water resources. This interpretation, however, is made cautiously because the ECM community of urban trees in water stress has not been studied. *Hebeloma sachariolens* was found only in park soil conditions where content of N and P was higher than in street soils, which is in agreement with the findings of many authors [56-59] regarding the ability of this fungus species to tolerate rather high nutrient conditions. The formation of mycorrhizae by *T. borchii* and *T. maculatum* is hardly surprising as the fungi have been reported to form mycorrhizal symbioses with *Tilia* spp. elsewhere in Europe [60,61].

Overall, our results showed that the tree vitality was significantly associated with soil characteristics, especially with heavy metal pollution. Our knowledge of ECM communities in urban areas is still limited, and these findings provide new insights into ECM distribution patterns in urban ecosystems. Given the multifunctional role of ECM in urban ecosystems, further research should also include manipulation of mycorrhizal communities in the field.

## Acknowledgements

We would like to thank the City Hall of Gdańsk and Road and Greenery Department of Gdańsk for founding of the data concerning the inventory of the Great Lime Avenue, which were the basis of the analyses carried out in this paper.

